# Antibody Resistance of SARS-CoV-2 Omicron BA.1, BA.1.1, BA.2 and BA.3 Sub-lineages

**DOI:** 10.1101/2022.04.07.487489

**Authors:** Jingwen Ai, Xun Wang, Xinyi He, Xiaoyu Zhao, Yi Zhang, Yuchao Jiang, Minghui Li, Yuchen Cui, Yanjia Chen, Rui Qiao, Lin Li, Lulu Yang, Yi Li, Zixin Hu, Wenhong Zhang, Pengfei Wang

## Abstract

The SARS-CoV-2 Omicron variant has been partitioned into four sub-lineages designated BA.1, BA.1.1, BA.2 and BA.3, with BA.2 becoming dominant worldwide recently by outcompeting BA.1 and BA.1.1. We and others have reported the striking antibody evasion of BA.1 and BA.2, but side-by-side comparison of susceptibility of all the major Omicron sub-lineages to vaccine-elicited or monoclonal antibody (mAb)-mediated neutralization are urgently needed. Using VSV-based pseudovirus, we found that sera from individuals vaccinated by two doses of inactivated whole-virion vaccines (BBIBP-CorV) showed very weak to no neutralization activity, while a homologous inactivated vaccine booster or a heterologous booster with protein subunit vaccine (ZF2001) markedly improved the neutralization titers against all Omicron variants. The comparison between sub-lineages indicated that BA.1.1, BA.2 and BA.3 had comparable or even greater antibody resistance than BA.1. We further evaluated the neutralization profile of a panel of 20 mAbs, including 10 already authorized or approved, against these Omicron sub-lineages as well as viruses with different Omicron spike single or combined mutations. Most mAbs lost their neutralizing activity completely or substantially, while some demonstrated distinct neutralization patterns among Omicron sub-lineages, reflecting their antigenic difference. Taken together, our results suggest all four Omicron sub-lineages threaten the efficacies of current vaccines and antibody therapeutics, highlighting the importance of vaccine boosters to combat the emerging SARS-CoV-2 variants.

## Introduction

The World Health Organization has now designated five variants of severe acute respiratory syndrome coronavirus 2 (SARS-CoV-2) as Variants of Concern, including Alpha (B.1.1.7), Beta (B.1.351), Gamma (P.1), Delta (B.1.617.2) and Omicron (B.1.1.529). The Omicron variant has recently been divided into four sub-lineages: BA.1, BA.1.1, BA.2 and BA.3 (Figure 1A). The original Omicron (BA.1 sub-lineage) was first identified in Botswana and South Africa in November 2021^1^ and together with its derivative BA.1.1 (containing an extra spike R346K mutation) became dominant worldwide in replacement of Delta over the span of a few weeks. But subsequently, we saw a rapid surge in the proportion of BA.2, and this sub-lineage became the dominant variant globally. Compared with the BA.1 and BA.2 sub-lineages, the prevalence of BA.3 sub-lineage is currently very low (Figure 1B).

**Figure 1.**
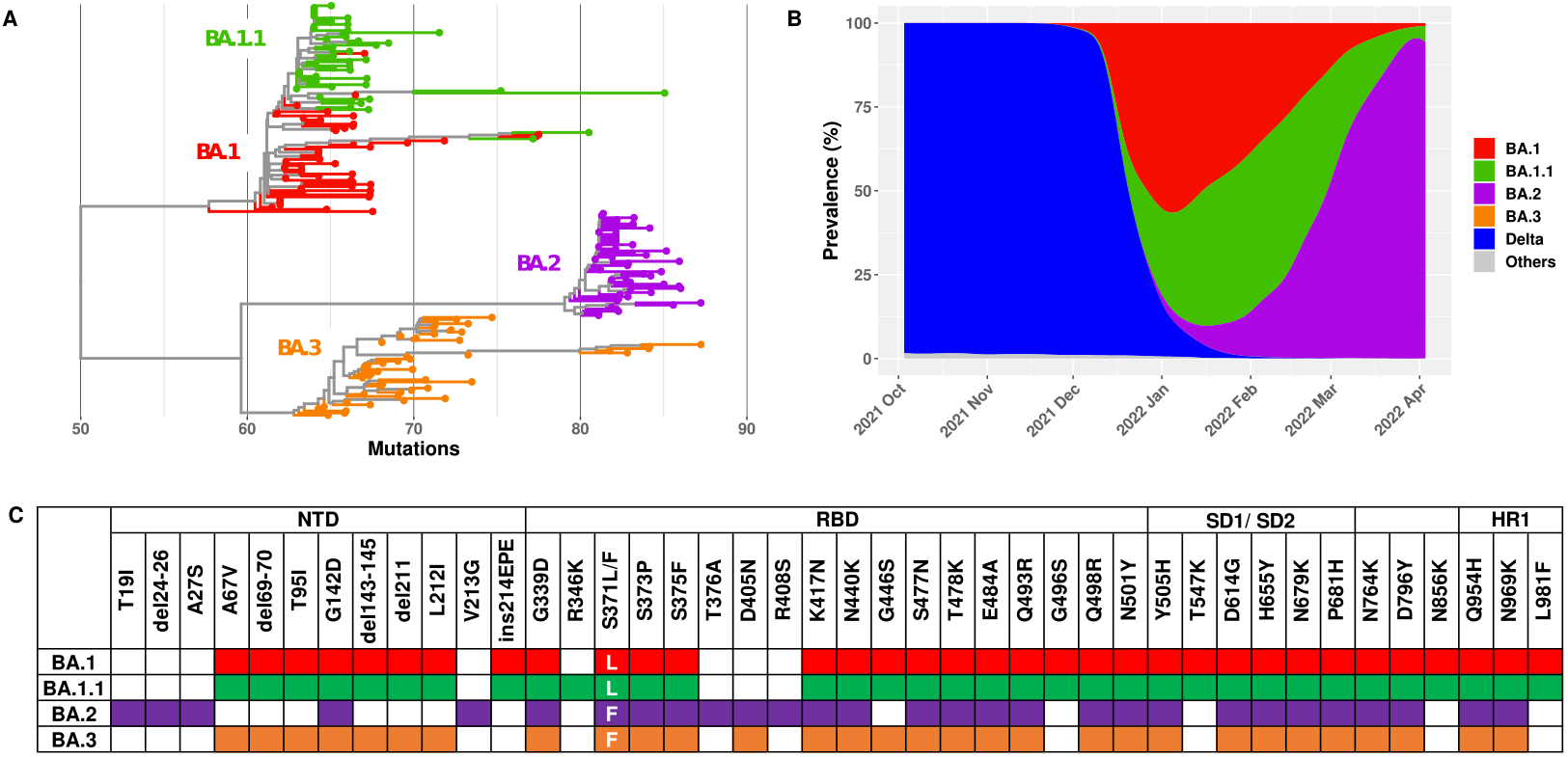
Characteristics of the Omicron sub-lineages. **(A)** Phylogenetic tree of the BA.1, BA.1.1, BA.2 and BA.3 sub-lineages. Fifty randomly selected sequences belonging to each of the Omicron sub-lineages from GISAID were used as query sequences **(B)** Prevalence of the Omicron sub-lineages and Delta variant based on all the sequences available on GISAID over the past six months. **(C)** Spike mutations within the Omicron sub-lineages.

BA.1, BA.2 and BA.3 have numerous mutations in common, but also with distinct sets of mutations in their spike that can differentiate these sub-lineages (Figure 1C). Although the selective advantage of BA.2 could be partially explained by its higher transmissibility than BA.1^2^, their relative immune evasion property could be also counted. We^3,4^ and others^5-12^ have reported that BA.1 demonstrated considerable escape from neutralization by monoclonal antibodies (mAbs) and sera from vaccinated individuals. BA.2 has also been reported to severely dampen antibody neutralization^13,14^. However, evaluation and comparison of susceptibility of all the major Omicron sub-lineages to vaccine-elicited or mAb-mediated neutralization are urgently needed. In this study, therefore, we constructed the Omicron sub-lineage pseudoviruses (PsVs) and compared side-by-side their neutralization sensitivity to vaccinee sera as well as a panel of mAbs.

## Results

The first question we asked for the Omicron sub-lineages is their extent of immune evasion of polyclonal antibody neutralizing activity elicited in humans after vaccination or infection. To answer this, we first assessed the neutralizing activity of sera from individuals vaccinated by two doses of inactivated whole-virion vaccines (BBIBP-CorV) (Supplementary Table 1). Similar as we reported before^3^, although all the sera showed neutralization activity against wild-type (WT) virus, the activity was relatively weak with geometric mean neutralizing titers (GMTs) about 55, and when it turned to BA.1, only 2/10 vaccinees showed marginal neutralization. When we tested these sera on the three other sub-lineages, most of them showed no detectable activity except a few had very weak neutralization against BA.2 and BA.3 (Figure 2A and Supplementary Figure 1A). These results indicate that two-dose inactivated vaccine is inadequate to provide full protection against these newly emerging Omicron variants. Our previous study showed that a booster shot, either homologous or heterologous, can reduce Omicron BA.1 escape from neutralizing antibodies^3^. To see if this is the case for the other Omicron sub-lineages, we then collected and tested 20 samples from healthy adults who had a third boosting vaccination shot with the same BBIBP-CorV vaccine (homologous booster group, Supplementary Table 1). As shown in Figure 2B, the sera had a neutralizing GMT against WT of 225 with 5- to 6-fold reduction against BA.1, BA.1.1, BA.2 and BA.3, but at least 15/20 samples exhibiting detectable neutralizing activity against all four sub-lineages. We also collected 18 sera from individuals that received two doses of BBIBP-CorV followed by a protein subunit vaccine (ZF2001) 4-8 months later (heterologous booster group, Supplementary Table 1). This cohort had higher neutralizing tiers with GMTs of 537, 108, 81, 42 and 69 against WT, BA.1, BA.1.1, BA.2 and BA.3, respectively. Although these numbers amount to 7- to 23-fold reductions of potency comparing Omicron sub-lineages to WT, almost all samples maintained detectable neutralizing activity against the Omicron variants (Figure 2C). The marked improvement in serum neutralization from individuals received a booster dose over those did not highlights the value of vaccine boosters for eliciting neutralizing antibody responses against Omicron sub-lineages. The emergence of the SARS-CoV-2 Delta variant led to an increasing number of breakthrough infection cases, to gain insight into their chance of re-infection by Omicron, we recruited 10 participants who were immunized with two-dose inactivated vaccines before infected by Delta variant (Supplementary Table 1). Serum samples were obtained from them after 3-4 months of breakthrough infection and evaluated on WT and the four Omicron sub-lineage PsVs (Figure 2D). We found that breakthrough infection by Delta boosted the neutralizing antibody titers significantly to very high levels against WT virus (GMT = 1,740). However, the neutralization titers for Omicron sub-lineages were significantly reduced, more than 100-fold in comparison to WT. The reduction level was much higher than that of the homologous and heterologous vaccine booster groups, which may be associated with the antigenic difference between Delta and Omicron variants.

**Figure 2.**
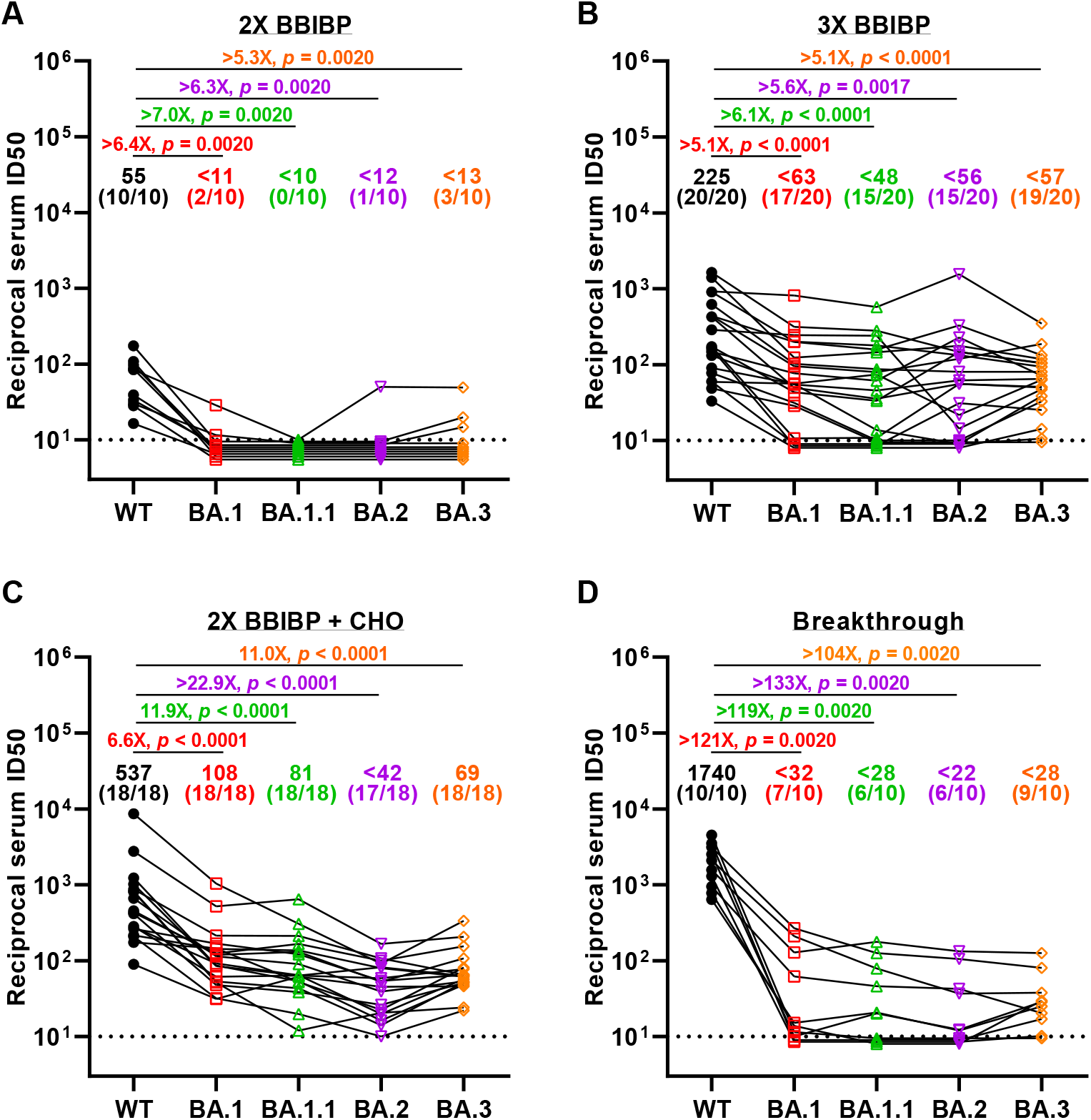
Neutralization of pseudotyped WT (D614G) and Omicron sub-lineage viruses by sera collected from individuals vaccinated with 2-dose BBIBP-CorV only (**A**), with a BBIBP-CorV homologous booster (**B**) or with a ZF001 heterologous booster dose (**C**) following two doses of BBIBP-CorV, or from individuals infected by Delta virus after vaccination (**D**). For all panels, values above the symbols denote geometric mean titer and the numbers in parentheses denote the proportion of positive sera with ID_50_ above the LOQ (dotted lines, >1:10). *P* values were determined by using a Wilcoxon matched-pairs signed-rank test (two-tailed).

Taking into account of all the serum samples, we also carried out a comparison between the original Omicron - BA.1 and the newly emerging sub-lineages to see if there are inherent difference regarding their immune evasion property. BA.1.1, with an additional R346K mutation on top of BA.1, showed slightly but statistically significant lower titers than BA.1. For BA.2 and BA.3, the neutralization titers were also lower than BA.1, which was mostly contributed by the heterologous booster group, indicating the receptor binding domain (RBD) subunit vaccine booster may induce some RBD-directed antibodies which could be evaded by the BA.2/BA.3 unique mutations (Supplementary Figure 2).

To better understand these differences and examine which types of antibodies in serum lose their activity against these Omicron sub-lineages, we further evaluated the neutralization profile of a panel of 20 mAbs targeting SARS-CoV-2 spike. These included 10 authorized or approved mAbs with sequences available: REGN10987 (imdevimab)^15^, REGN10933 (casirivimab)^15^, LY-CoV555 (bamlanivimab)^16^, CB6/LY-CoV016 (etesevimab)^17^, S309 (sotrovimab)^18^, COV2-2130 (cilgavimab)^19^, COV2-2196 (tixagevimab)^19^, CT-P59 (regdanvimab)^20^, Brii-196 (amubarvimab)^21^ and LY-CoV1404 (bebtelovimab)^22^, all of which are directed to RBD. We also included some other RBD-directed mAbs of interest, including 1-20, 2-15, 1-57, 2-7^23^, and 2-36^23,24^ from our own collection and ADG-2^25^ from Adagio Therapeutics, and four more NTD-directed mAbs, including 5-24, 4-18, 4-19^23,26^ and 5-7^23,27^. Overall, all four Omicron sub-lineages had severe impact on most of the antibodies but they also showed important differences in neutralization patterns. Among the authorized or approved mAbs, seven were either totally inactive or severely impaired in neutralizing all four sub-lineages. S309, the only approved antibody found to retain its neutralizing activity against the original form of Omicron in our previous study^3^, lost more neutralizing activity against BA.2 and BA.3. COV2-2130 completely lost its neutralizing activity against BA.1 and BA.1.1, while retaining largely active against BA.2 and BA.3. Luckily, LY-CoV1404, which has been granted emergency use authorization very recently, remained potent in neutralizing all Omicron sub-lineages, continuing to broaden its coverage of SARS-CoV-2 variants^28^. For the other RBD- or NTD-directed mAbs, none of them retained full neutralizing activity against all of the Omicron sub-lineages. Two class 4 antibodies, ADG-2 and 2-36, retained decent activity against BA.1 and BA.1.1 but lost their neutralizing activity completely against BA.2 and BA.3. Interestingly, 2-7, one of our class 3 antibody, completely lost its neutralizing activity against BA.1, BA.1.1 and BA.3, while retaining largely active against BA.2. Similar pattern was seen for another approved class 3 antibody, REGN10987. On the contrary, we found the activity of 5-7, the non-supersite-directed NTD antibody, was partially retained against BA.1, BA.1.1 and BA.3, but totally abolished against BA.2 (Figure 3A and Supplementary Figure 3).

**Figure 3.**
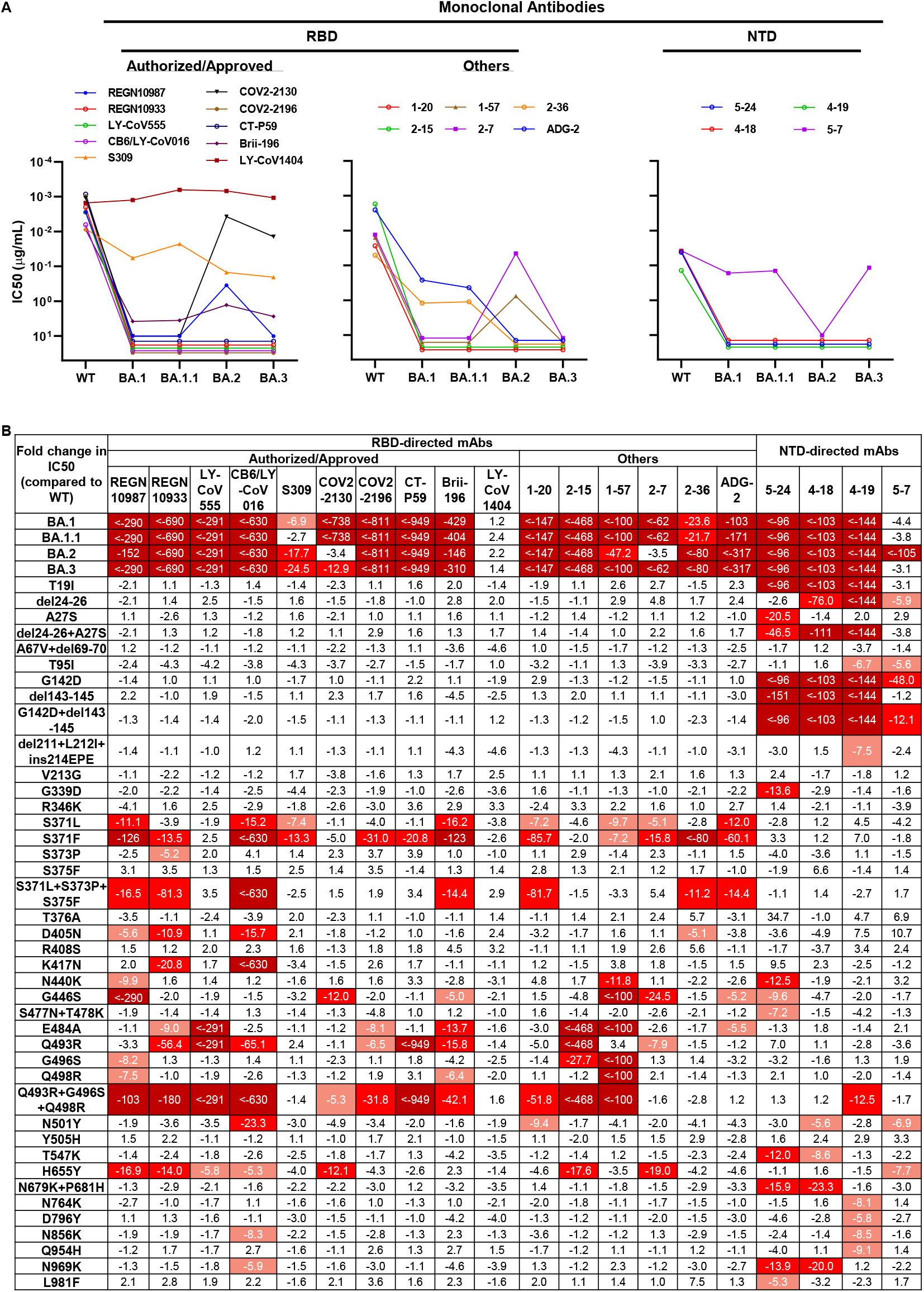
Neutralization of pseudotyped WT (D614G) and Omicron sub-lineage viruses by mAbs targeting different epitopes. **(A)** Changes in neutralization IC_50_ of select RBD and NTD mAbs against Omicron sub-lineage pseudoviruses. **(B)** Fold increase or decrease in neutralization IC_50_ of mAbs against Omicron sub-lineage and single-mutation as well as combined-mutation pseudoviruses, relative to WT, presented as a heatmap with darker colors implying greater change.

To dissect the key mutations conferring antibody resistance and the specific mutations leading to the different neutralization patterns of Omicron sub-lineages, we constructed PsVs with each of the spike single mutations alone or in combination if they are spatially close and tested them using the same panel of 20 mAbs. Totally 40 specific mutation viruses were tested and their comprehensive neutralization profile by these 20 mAbs are summarized in Figure 3B as fold change in 50% inhibitory concentration (IC_50_) relative to WT virus. For mutations affecting antibody activity, the first ones caught our attention were S371L and S371F, both broadly affected most of the RBD-directed mAbs with S371F had a greater negative impact. Intriguingly, when we tested S371L, S373P and S375F in combination, as they form a loop adjacent to a lipid-binding pocket^29^, we indeed observed synergistic effect of the triple serine mutations in the reduction of neutralization potency of some mAbs. Q493R appears to be another key mutation responsible for the loss in potency of many RBD antibodies, and again, when it was tested in combination with G496S and Q498R, apparent synergistic effect was seen for some mAbs. G446S, which is lacking in BA.2 but presented in the other sub-lineages, may explain why COV2-2130 and 2-7 are not much affected by BA.2. Other mutations, such as D405N, K417N, N440K, E484A and N501Y, distinctly affected the activity of different RBD-directed mAbs, most of which could be explained by the mutations falling into the antibody epitopes. For LY-CoV1404, same as we saw for the Omicron sub-lineages, none of the single mutations significantly affected its neutralization potency, indicating despite of the constellation of spike mutations present in these viruses, there is still a patch within LY-CoV1404’s binding region that is not affected. For the NTD-directed mAbs, it was mostly the mutations falling into the NTD of the spike, including T19I, del24-26+A27S and G142D+del142-145, that are responsible for the neutralization activity lost as expected.

## Discussion

The SARS-CoV-2 Omicron variant immediately raised alarms after its identification and the scenario seems to getting worse with the emerging Omicron sub-lineages, like BA.2, which has been reported to be inherently substantially more transmissible than BA.1^2^. Lots of research articles have been published studying the original Omicron BA.1 virus, but less is known about the BA.2 and other sub-lineages. Here in this study, we constructed all the major Omicron sub-lineage viruses to date - BA.1, BA.1.1, BA.2 and BA.3, and investigated their antibody evasion property in parallel.

We previously reported the markedly reduced neutralizing activity against BA.1 of convalescent or BBIBP-CorV vaccination sera^3,4^. Here, we showed all polyclonal sera also had a substantial loss in neutralizing activity against the other Omicron sub-lineages, with drops comparable to or even greater than that of BA.1, indicating all these sub-lineages have a very far antigenic distance from the WT virus. Our results are quite comparable to studies on the mRNA vaccines^13,30^, showing neutralizing antibody titer against BA.2 was similar to or lower than that against BA.1. Based on these, we suggest the selective advantage of BA.2 over BA.1 should be mainly contributed by its higher transmissibility rather than immune evasion. On the other hand, we showed a third homologous inactivated vaccine booster or a heterologous protein subunit vaccine booster could elevate neutralization titer against BA.1 significantly^3^. This is also true for the other Omicron sub-lineages. Most recently, three recombinant lineages (XD, XE and XF) have been reported^31^, but their antibody evasion should not be significantly different from the Omicron sub-lineages studied here since their spikes are identical to either BA.1 or BA.2. Therefore, promotion and popularization of vaccine booster injection is still an effective means to prevent SARS-CoV-2 transmission.

We also investigated the immune evasion capacity of Omicron sub-lineages with mAbs. Similar as we reported for BA.1^3^, most mAbs lost their neutralizing activity against BA.1.1, BA.2 and BA.3 completely or substantially. But we did observe some distinct neutralization patterns for certain mAbs among these sub-lineages, reflecting their different mutations. For example, S309 and 5-7, targeting some unique sites in RBD^18^ or NTD^27^, were the two mAbs reported to retain largely activity against BA.1^3,23^, but their activity was further abolished by BA.2. On the contrary, some mAbs like COV2-2130 and 2-7 lost activity against BA.1 totally but regained activity against BA.2. The good news is LY-CoV1404 or bebtelovimab kept its potent neutralization activity against all Omicron sub-lineages and other major SARS-CoV-2 variants^28,32^, standing out like a lone star in the darkness. Our data are in good consistency with others^5,13^ regarding the mAb neutralization profile of the Omicron sub-lineages and single mutations, but we had more sub-lineage - BA.3, and combined some mutations in proximity to investigate their synergistic actions.

Although LY-CoV1404 remains to be our hope of SARS-CoV-2 therapeutic antibodies currently, resistance may arise sometime if it is administered as mono-therapy for a prolonged period given the error-prone property of RNA virus. Therefore, it is advisable to develop more potent and broad neutralizing antibodies to be administered as cocktail to contain this ever-evolving pathogen. Meanwhile, vaccine boosters, either homologous or heterologous, could elicit neutralizing antibodies that help reduce the viral escape and should be push forward.

## Methods

### Serum samples

Sera from individuals who received two or three doses of BBIBP-CorV or ZF2001 vaccine were collected at Huashan Hospital, Fudan University 14 days after the final dose. Sera were also obtained from patients after 3-4 months of SARS-CoV-2 breakthrough infection caused by Delta variant after immunizing with two-dose inactivated vaccines (CoronaVac). All collections were conducted according to the guidelines of the Declaration of Helsinki and approved by the Institutional Review Board of the Ethics Committee of Huashan Hospital (2021-041 and 2021-749). All the participants provided written informed consents.

### Monoclonal antibodies

Monoclonal antibodies tested in this study were constructed and produced at Fudan University.

### Construction and production of variant pseudoviruses

Plasmids encoding the WT (D614G) SARS-CoV-2 spike and Omicron sub-lineage spikes, as well as the spikes with single or combined mutations were synthesized. Expi293 cells were grown to 3×10^6^/mL before transfection with the indicated spike gene using Polyethylenimine (Polyscience). Cells were cultured overnight at 37°C with 8% CO_2_ and VSV-G pseudo-typed ΔG-luciferase (G*ΔG-luciferase, Kerafast) was used to infect the cells in DMEM at a multiplicity of infection of 5 for 4 h before washing the cells with 1×DPBS three times. The next day, the transfection supernatant was collected and clarified by centrifugation at 300g for 10 min. Each viral stock was then incubated with 20% I1 hybridoma (anti-VSV-G; ATCC, CRL-2700) supernatant for 1 h at 37 °C to neutralize the contaminating VSV-G pseudotyped ΔG-luciferase virus before measuring titers and making aliquots to be stored at −80 °C.

### Pseudovirus neutralization assays

Neutralization assays were performed by incubating pseudoviruses with serial dilutions of monoclonal antibodies or sera, and scored by the reduction in luciferase gene expression. In brief, Vero E6 cells were seeded in a 96-well plate at a concentration of 2×10^4^ cells per well. Pseudoviruses were incubated the next day with serial dilutions of the test samples in triplicate for 30 min at 37 °C. The mixture was added to cultured cells and incubated for an additional 24 h. The luminescence was measured by Luciferase Assay System (Beyotime). IC_50_ was defined as the dilution at which the relative light units were reduced by 50% compared with the virus control wells (virus + cells) after subtraction of the background in the control groups with cells only. The IC_50_ values were calculated using nonlinear regression in GraphPad Prism.

### Sequence alignment and phylogenetic tree

This analysis involved 200 nucleotide sequences, including 50 samples for each lineage randomly selected from the GISAID database. Sequence alignment was carried out using ClustalW progress^33^ and corrected manually. The evolutionary history was inferred using the Neighbor-Joining method^34^. The optimal tree is shown. The tree is drawn to scale, with branch lengths in the same units as those of the evolutionary distances used to infer the phylogenetic tree. The evolutionary distances were computed using the p-distance method^35^ and are in the units of the number of base differences per site. All positions with less than 50% site coverage were eliminated. There was a total of 29743 positions in the final dataset. Evolutionary analyses were conducted in MEGA11^36^ and visualized with the package ‘ggtree’ in R. The current snapshot of COVID-19 data was taken from GISAID between Oct 2021 and Mar 2022 in weekly basis. Lineage level prevalence rate was summarized using cubic spline interpolation.

## Data availability

Materials used in this study will be made available but may require execution of a materials transfer agreement. All the data are provided in the paper or the Supplementary Information.

## Acknowledgements

This research was supported by Shanghai Municipal Science and Technology Major Project (2017SHZDZX01).

## Author contributions

P.W., W.Z., and Z.H. conceived and supervised the project. J.A., X.W., X.Z., Y.Z., Y.J., M.L., Y.Cui, Y.Chen, R.Q., L.L., and L.Y. conducted the biological experiments. X.H., Y.L., and Z.H. conducted the bioinformatics analysis. J.A., X.W., X.Z., Y.Z., Y.J., Z.H., W.Z., and P.W. analyzed the results and wrote the manuscript. All the authors reviewed, commented, and approved the manuscript.

## Supplementary Figures

**Supplementary Figure 1.**
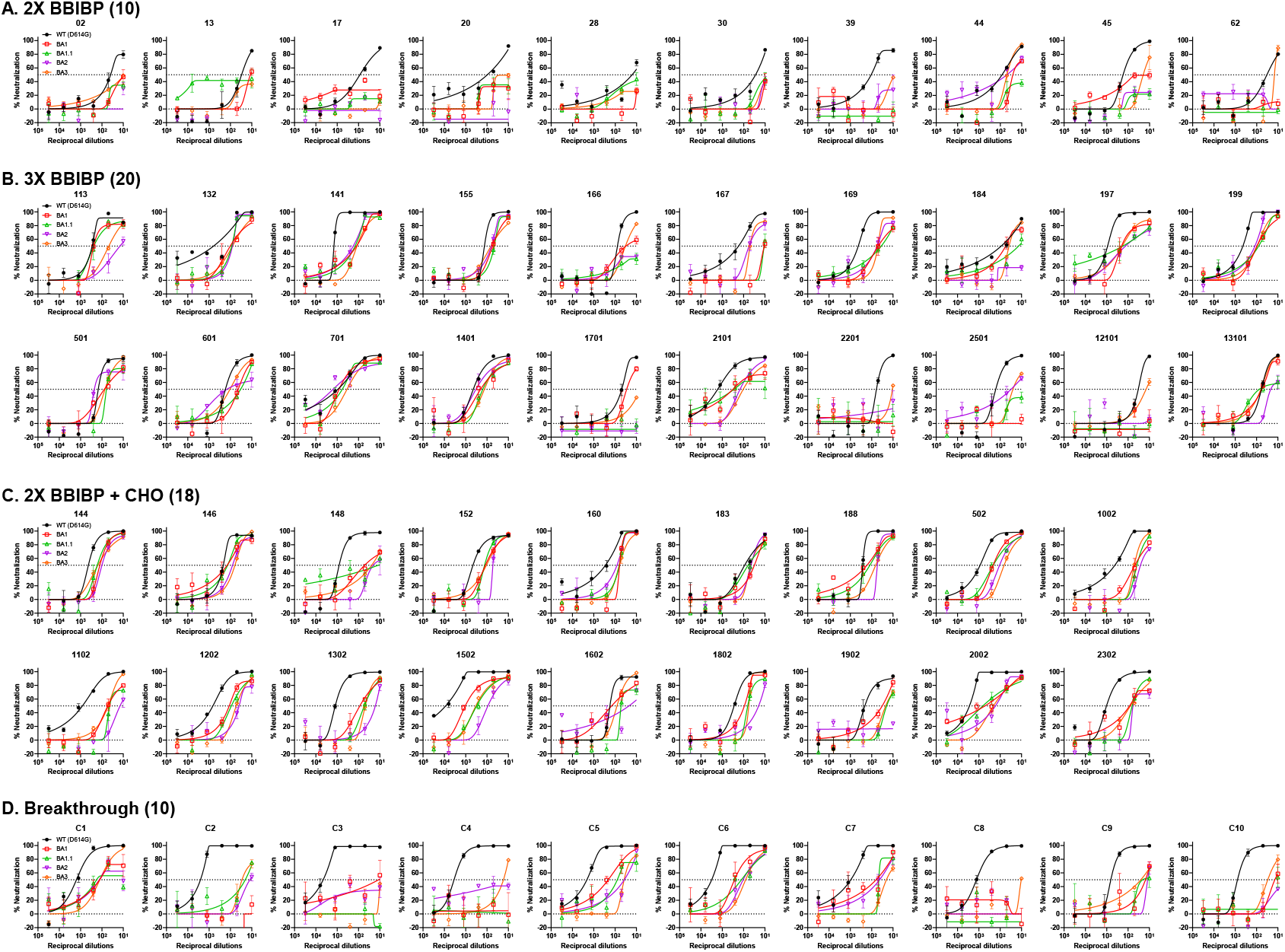
Neutralization curves for sera collected from individuals vaccinated with 2-dose BBIBP-CorV only (**A**), with a BBIBP-CorV homologous booster (**B**) or with a ZF001 heterologous booster dose (**C**) following two doses of BBIBP-CorV, or from individuals infected by Delta virus after vaccination (**D**).

**Supplementary Figure 2.**
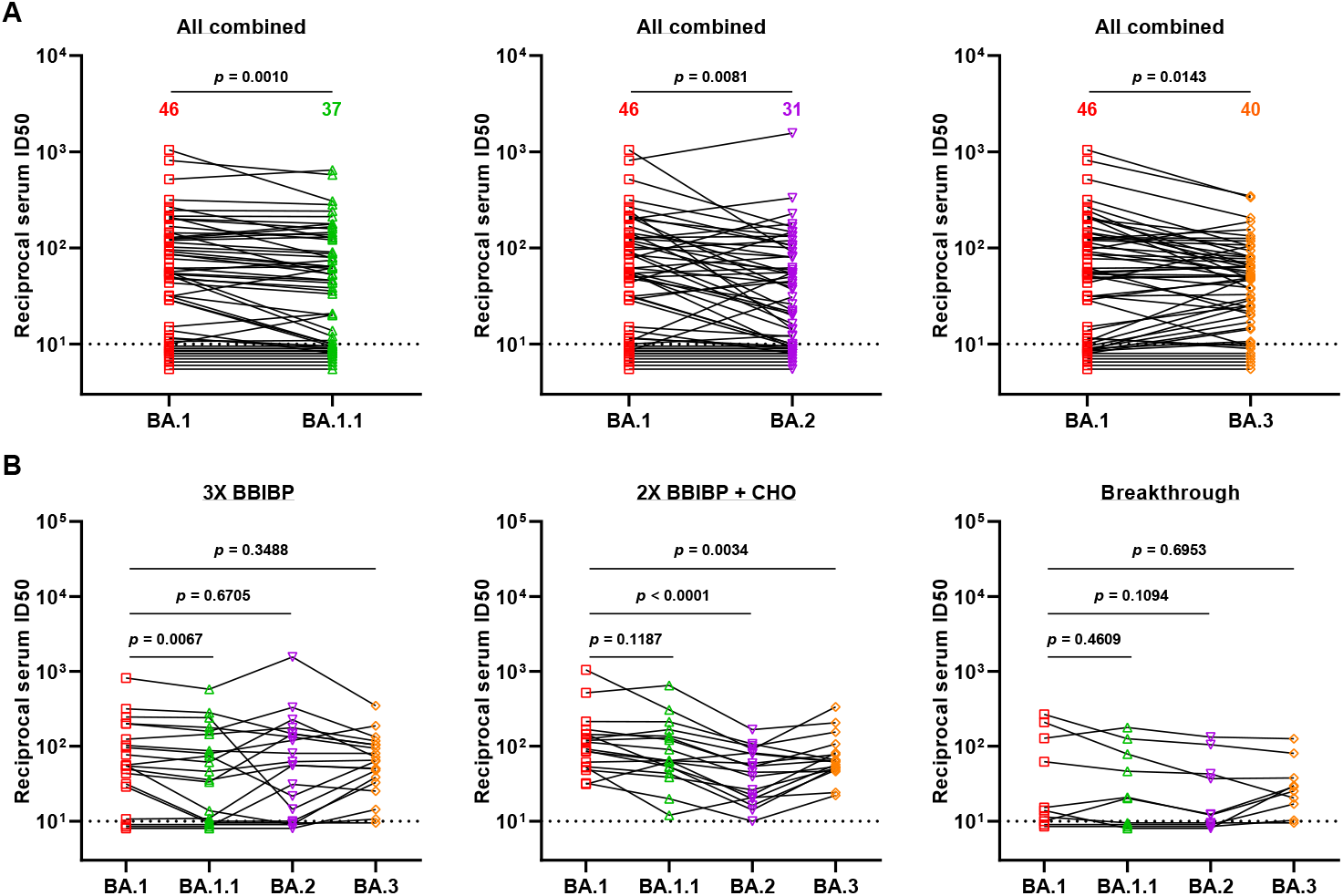
Comparison between BA.1 and the other Omicron sub-lineages with all the sera neutralization data combined (**A**) or within the different immunization groups (**B**). *P* values were determined by using a Wilcoxon matched-pairs signed-rank test (two-tailed).

**Supplementary Figure 3.**
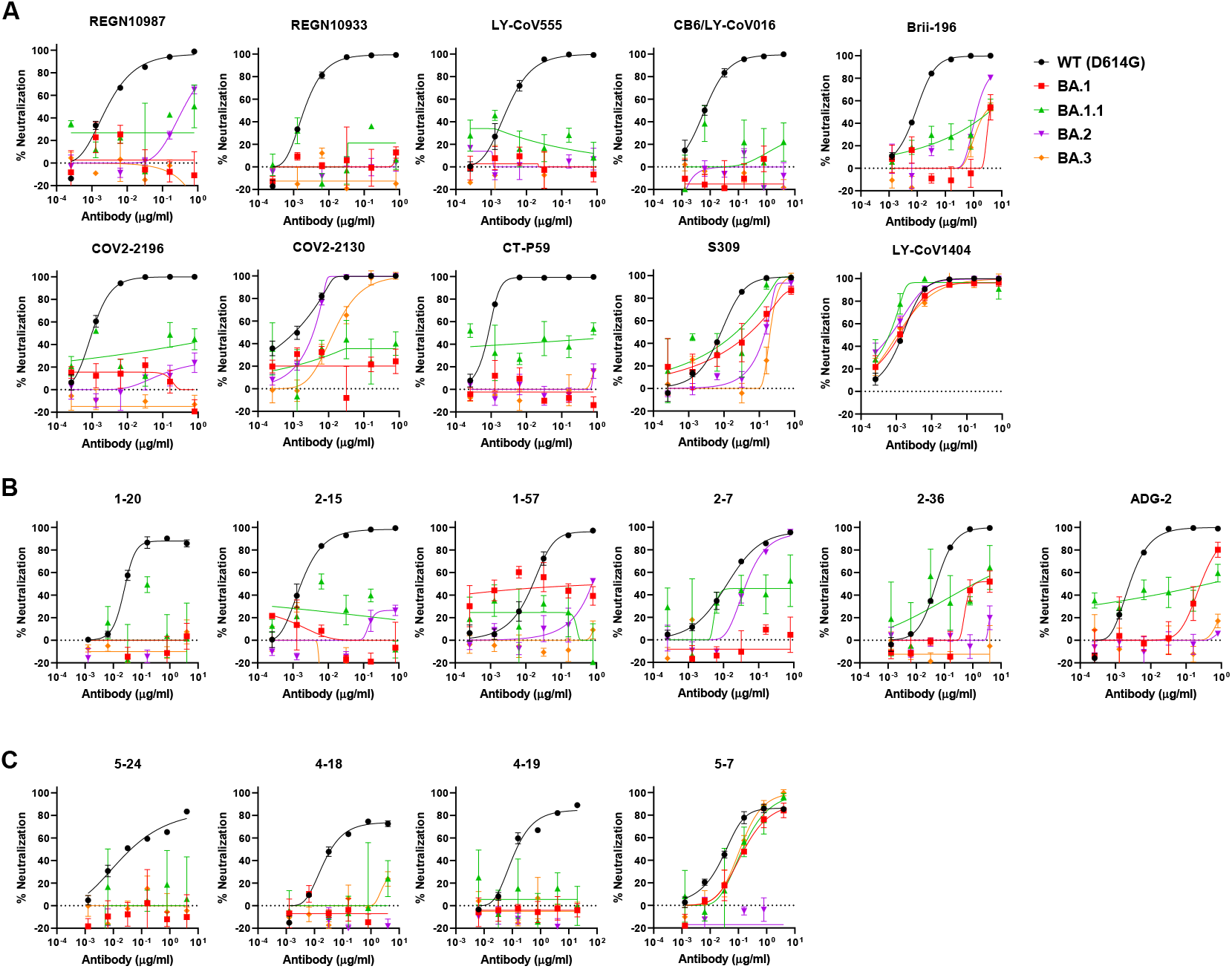
Neutralization curves for mAbs against WT (D614G) and Omicron sub-lineage viruses.

**Supplementary Table 1.** Baseline characteristics of enrolled participants, including Breakthrough infection group, BBIBP-CorV two doses group, BBIBP-CorV homologous booster group and BBIBP-CorV/ZF2001 heterologous booster group.

|  | Breakthrough<br>infection<br>(n=10) | BBIBP-CorV<br>two doses<br>(n=10) | BBIBP-CorV<br>homologous<br>booster<br>(n=20) | BBIBP-CorV/<br>ZF2001<br>heterologous<br>booster<br>(n=18) | P value |
| --- | --- | --- | --- | --- | --- |
| **Age (years),**<br>**median(range)** | 46 (34-54) | 31.5 (23-51) | 28 (21-59) | 29.5 (23-53) | < 0.001 |
| **Male, n (%)** | 3 (30.00%) | 2 (20.00%) | 8 (40.00%) | 4 (22.22%) | 0.581 |
| **BMI (kg/m<sup>2</sup>), mean (SD)** | 24.45(5.64) | 23.35 (3/16) | 22.25 (2.96) | 20.56 (2.01) | 0.028 |
| **Comorbidities (%)** |  |  |  |  |  |
| Any, n (%) | 4 (40.00%) | 0 (0.00%) | 2 (10.00%) | 0 (0.00%) | 0.008 |
| Cardiovascular diseases,<br>n (%) | 0 (0.00%) | 0 (0.00%) | 0 (0.00%) | 0 (0.00%) | / |
| Hypertension, n (%) | 3 (30.00%) | 0 (0.00%) | 0 (0.00%) | 0 (0.00%) | 0.008 |
| Diabetes, n (%) | 1 (10.00%) | 0 (0.00%) | 0 (0.00%) | 0 (0.00%) | 0.345 |

